# Atypical Chemokine Receptor DARC/ACKR1 Is Expressed Selectively by Neurons with Regional Heterogeneity and Protects against Neuroinflammation and Cognitive Impairment

**DOI:** 10.64898/2026.09.08.750199

**Authors:** V. N. Sankeerth Kanumuri, Ming Guo, Jayvon Nougaisse, Xujun Ma, Quansheng Du, Neal L. Weintraub, Yun Lei, Xin-Yun Lu

**Affiliations:** Department of Neuroscience & Regenerative Medicine, Medical College of Georgia at Augusta University, GA, USA; Department of Medicine, Medical College of Georgia at Augusta University, GA, USA; Vascular Biology Center, Medical College of Georgia at Augusta University, GA, USA

## Abstract

Atypical chemokine receptors (ACKRs) regulate inflammatory responses by scavenging and sequestering chemokines. The Duffy antigen receptor for chemokines (DARC), also known as ACKR1, binds multiple inflammatory chemokines and regulates inflammation in peripheral tissues. However, the distribution and functional role of DARC/ACKR1 in the brain remain to be defined. Here, we show that DARC/ACKR1 is widely expressed throughout the mouse brain, with marked regional heterogeneity. DARC/ACKR1 is predominantly expressed by neurons and is not detected in microglia or astrocytes. Importantly, this neuronal expression pattern was also observed in the human brain. Furthermore, DARC/ACKR1 deficiency resulted in enhanced neuroinflammation, characterized by increased microglial activation and astrogliosis across multiple brain regions, particularly the hippocampus, and was accompanied by cognitive deficits. Together, these findings identify DARC/ACKR1 as a previously unrecognized neuron-specific atypical chemokine receptor that plays an important role in restraining neuroinflammation and preserving cognitive function.

## INTRODUCTION

Chemokines are a large family of small, secreted proteins that direct cell migration and positioning throughout the body, primarily targeting cells involved in inflammation. Structurally, chemokines are classified into C, CC, CXC, and CX3C subfamilies based on the number and spacing of their cysteine residues in their N terminus, where “C” denotes cysteine and “X” represents any amino acid residue (1, 2). Functionally, chemokines are broadly classified as homeostatic or inflammatory (3, 4). Homeostatic chemokines are constitutively expressed and regulate immune surveillance, immune cell positioning and tissue organization during steady-state conditions. In contrast, inflammatory chemokines are induced in response to inflammatory stimuli and function to recruit and activate immune cells, thereby amplifying and sustaining inflammatory responses that can become detrimental when prolonged or dysregulated.

Chemokines bind to two major classes of chemokine receptors on target cells: classical chemokine receptors, which are members of the G protein-coupled receptor (GPCR) superfamily and primarily mediate G protein-dependent signaling that regulates chemotaxis and immune cell activation, and atypical chemokine receptors (ACKRs), which are structurally related to classical chemokine receptors but lack canonical G protein-mediated signaling capacity and instead regulate chemokine availability through ligand scavenging, transport, and sequestration (5–7). A key feature of the chemokine system is its extensive receptor–ligand promiscuity, whereby a single chemokine can bind to multiple receptors and a single receptor can recognize multiple chemokines, resulting in substantial functional redundancy, particularly among inflammatory chemokines (5, 8). Despite this promiscuity, functional specificity of the chemokine system is achieved through the selective expression of canonical receptors across distinct cell types and tissues, while ACKRs contribute an additional layer of functional specificity by regulating the availability and spatial distribution of chemokines, thereby controlling their access to target cells to modulate chemokine-driven inflammatory responses.

The Duffy antigen receptor for chemokines (DARC), also known as atypical chemokine receptor 1 (ACKR1), was first identified in the 1950s as a red blood cell (RBC) blood-group antigen and named after the patient in whom it was discovered (9, 10). It was subsequently recognized as an erythrocyte receptor exploited by *Plasmodium vivax* for invasion (11) and was later identified as a silent non-signaling chemokine receptor that binds highly promiscuously to inflammatory CC and CXC chemokines (12–14). Through ligand scavenging and sequestration, DARC/ACKR1 regulates the availability and distribution of these chemokines and can limit excessive inflammatory responses and promote the resolution of inflammation in peripheral tissues.

Although DARC/ACKR1-binding CC and CXC chemokines are expressed in the central nervous system (CNS), and their canonical receptors have been extensively studied in neuroinflammation, little is known about the expression and functional role of DARC/ACKR1 in the brain (15). To date, only two earlier studies have described DARC/ACKR1 expression in the cerebellum (16, 17). Its cellular and regional distribution throughout the brain, as well as its function, remain to be defined. Given that several DARC/ACKR1-binding chemokines (18), including CCL2/MCP-1 and CCL5 (19–21) as well as CXCL1 (22) and CXCL5 (23, 24), are established mediators of neuroinflammation, we hypothesized that DARC/ACKR1 in the brain may contribute to the regulation of neuroinflammatory processes by controlling inflammatory chemokine availability. In the present study, we first characterized the regional and cellular distribution of DARC/ACKR1 in the brain. We then examined microglial activation, astrogliosis, and cognitive function in DARC/ACKR1 knockout mice to determine the impact of DARC/ACKR1 loss on neuroinflammation and cognitive function.

## MATERIALS AND METHODS

### Human postmortem brain tissues

Human postmortem brain tissues were obtained from the Neuropathology Core of the Emory Alzheimer’s Disease Research Center. All specimens were fully de-identified. Accordingly, this study was reviewed by the Augusta University Institutional Review Board (IRB) and determined not to constitute human subjects research.

### Animals

Male and female C57BL/6J mice were obtained from The Jackson Laboratory (Bar Harbor, MA, USA) and maintained in our colony. C57BL/6J mice were used for regional and cellular mapping of DARC/ACKR1 expression in the brain. DARC/ACKR1 knockout mice on a C57BL/6J background, carrying a targeted deletion in the 5′ region of exon 2 of the mouse *Ackr1* gene (25), were originally obtained from the Mutant Mouse Resource and Research Centers (MMRRC-UNC, strain no. 029873; University of North Carolina, Chapel Hill, NC). Heterozygous male and female mice were intercrossed to generate homozygous DARC/ACKR1 knockout (DARC KO) mice and wild-type (WT) littermate controls. Mice were group housed in individually ventilated cages, with *ad libitum* access to food and water under a 12 h light–dark cycle (lights on at 6:00 AM). All procedures were conducted in accordance with the National Institutes of Health (NIH) Guide for the Care and Use of Laboratory Animals and approved by the Institutional Animal Care and Use Committee of Augusta University.

### Immunofluorescence staining

Mice were transcardially perfused under anesthesia with 0.1 M phosphate-buffered saline (PBS), followed by 4% paraformaldehyde (PFA) in PBS. Brains were post-fixed overnight in 4% PFA at 4°C and then cryoprotected in 30% sucrose in PBS until fully equilibrated. Coronal brain sections (40 µm thick) were cut using a cryostat and collected as free-floating sections. Sections were stored in cryoprotectant solution at −20°C until further processing. Free-floating brain sections were rinsed in PBS and incubated in blocking solution containing 10% donkey serum and 0.5% Triton X-100 in Tris-buffered saline (TBS) for 1.5 h at room temperature and then incubated overnight at 4°C with primary antibodies against DARC/ACKR1 (LSBio, LS-B2306, 1:250), neuronal nuclear antigen (NeuN; Sigma-Aldrich, ABN90P, 1:500), ionized calcium-binding adapter molecule 1 (IBA1; Wako/Fujifilm, 019-19741, 1:500), glial fibrillary acidic protein (GFAP; Cell Signaling Technology, 3670, 1:500), glutamate decarboxylase 67 (GAD67; Millipore, MAB5406, 1:500), and monocyte chemoattractant protein-1/C-C motif chemokine ligand 2 (MCP-1/CCL2; Novus Biologicals, NBP1-07035, 1:500). Following primary antibody incubation, sections were washed four times in TBS for 10 min each and incubated with the corresponding species-specific Alexa Fluor–conjugated secondary antibodies (1:1000) for 2 h at room temperature. Sections were subsequently washed in TBS and counterstained with 4′,6-diamidino-2-phenylindole (DAPI) to visualize cell nuclei. After the final washes, sections were mounted onto glass slides using Immuno-Mount and cover slipped for fluorescence imaging.

Frozen human postmortem brain tissues from the dorsolateral prefrontal cortex (DLPFC; Brodmann area 9) of neurologically normal control subjects were cut and stored at −80°C until use. Tissue sections were fixed in 4% PFA for 15 min at room temperature and then washed three times in TBS for 10 min each. Sections were blocked for 1 h at room temperature in TBS containing 0.5% (v/v) Triton X-100 and 10% donkey serum, followed by overnight incubation at 4°C with anti-DARC/ACKR1 and anti-NeuN antibodies diluted in the same blocking solution. The following day, sections were washed four times in TBS for 10 min each and incubated for 2 h at room temperature with the Alexa Fluor-conjugated secondary antibodies diluted in TBS containing 0.5% (v/v) Triton X-100. Sections were briefly rinsed in TBS and counterstained with DAPI to visualize nuclei, followed by three additional washes in TBS for 10 min each. Brain tissue sections were cover-slipped with an antifade mounting medium. Fluorescence images were acquired using the imaging procedures described above.

### RNAscope *in situ* hybridization

mRNA expression of *Ackr1* (DARC) and *Rbfox3* (NeuN) was detected using the RNAscope Multiplex Fluorescent V2 assay (Advanced Cell Diagnostics, Newark, CA, USA) in fresh-frozen brain sections. Mice were euthanized by cervical dislocation and rapidly decapitated. Brains were dissected on ice and snap-frozen by repeated immersion in isopentane (2-methylbutane) cooled to −30 to −40°C on dry ice. Coronal sections (16 µm thick) were cut on a cryostat, thaw-mounted onto SuperFrost Plus slides (Thermo Fisher Scientific), air-dried for 1 h at room temperature, and stored at −80°C until use. Briefly, brain sections were fixed in cold 4% paraformaldehyde for 60 min at 4°C, rinsed in PBS, and dehydrated through a graded ethanol series (50%, 70%, 100%, and 100%; 5 min each) at room temperature. Following rehydration in double-distilled water (ddH₂O), endogenous peroxidase activity was quenched with hydrogen peroxide for 10 min at room temperature. Sections were then subjected to target retrieval in ACD Retrieval Buffer for 5 min at 99–102°C, rinsed in ddH₂O, dehydrated in 100% ethanol for 3 min, and treated with Pretreat Pro Protease for 30 min at room temperature. Tissue was kept hydrated throughout the pretreatment procedure. RNAscope HiPlex probes targeting mouse *Ackr1*(ACD catalog no. 450231; Advanced Cell Diagnostics) and *Rbfox3* (ACD catalog no. 313311; Advanced Cell Diagnostics) were hybridized for 2 h at 40°C in a HybEZ oven. Signal amplification was performed according to the manufacturer’s protocol using AMP1 (30 min), AMP2 (30 min), and AMP3 (15 min), all at 40°C, with two 2-min washes in ACD Wash Buffer between amplification steps. For multiplex detection, channel-specific horseradish peroxidase (HRP) adapters (HRP-C1, HRP-C2, and HRP-C3; 15 min each at 40°C) were sequentially applied, each followed by Opal fluorophore deposition (1:1500 in TSA buffer; 30 min at 40°C) and HRP blocking (15 min at 40°C), with Wash Buffer rinses between all incubations. Sections were counterstained with DAPI for 30 sec and cover-slipped with ProLong Gold Antifade Mountant (Thermo Fisher Scientific).

### Image acquisition and quantification

For whole-section overview imaging, a KEYENCE BZ-X800 All-in-One Fluorescence Microscope (Keyence Corporation of America, Itasca, IL, USA) was used to visualize immunostaining and acquire images. All other fluorescence imaging was performed using a Leica STELLARIS 5 laser-scanning confocal microscope (Leica Microsystems, Wetzlar, Germany) equipped with a 405-nm diode laser, a tunable white-light laser (WLL; 440– 790 nm), and Power HyD S detectors. The system was operated using LAS X software with the Navigator module for automated tile scanning (Leica Microsystems). Regional overviews of selected brain structures were acquired using a 10× objective as mosaic tile scans with z-stack acquisition. Higher-magnification z-stack images were acquired using a 20× objective. Sequential scanning was used for multichannel image acquisition. Within each immunostaining experiment, the z-step size, laser power, detector gain, and pinhole settings were kept constant across all samples. For RNAscope experiments, imaging parameters were optimized according to the emission spectra of the Opal fluorophores, and mosaic tile scans were acquired using the same imaging workflow. All images were exported as TIFF files and processed using Fiji (ImageJ).

### Microglia quantification

Microglial density and soma morphology were assessed from IBA1-immunolabeled sections following established ImageJ-based quantification protocols (26). Confocal z-stacks acquired at 20× were converted to maximum-intensity projections using Fiji. For each section, regions of interest (ROI) were manually delineated. ROI areas (mm²) were recorded for normalization. IBA1+ microglia within each ROI were counted manually using the Cell Counter plugin in Fiji. Only cells with a clearly identifiable soma falling entirely within the ROI boundary were scored; cells intersecting the exclusion edges were omitted to prevent double counting across adjacent fields. Microglial density was expressed as IBA1+ cells per mm² of ROI area. Microglial soma area was quantified as a morphological index of activation state, as reactive microglia undergo somatic hypertrophy concurrent with process retraction (26, 27). Maximum-intensity projections of the IBA1 channel were converted to 8-bit grayscale and processed using a fast Fourier transform (FFT)-based bandpass filter to suppress low-frequency background variations and high-frequency image noise while preserving spatial features of interest. Image contrast was enhanced using the unsharp mask function, and the despeckle filter was applied to remove residual noise. A consistent automated threshold (Otsu method) was applied to generate binary masks of IBA1-positive signal. To isolate microglial somata from processes, a morphological opening filter (1-pixel-radius octagonal structuring element) was applied to the binary image. Individual soma areas (µm²) were measured using the analyze particles function with minimum and maximum particle size thresholds applied to exclude sub-cellular debris and merged soma clusters, respectively. Mean soma area was calculated per ROI, averaged across sections, and then across animals for group comparisons.

### Astrocyte quantification

Astrocytic reactivity was evaluated from GFAP-immunolabeled sections using two complementary measures: GFAP+ percent area coverage and individual astrocyte cell area (28). Percent area coverage was used as a global index of astrogliosis that captures both upregulation of GFAP expression and hypertrophic expansion of astrocytic processes. Maximum-intensity z-projections of the GFAP channel were converted to 8-bit grayscale in Fiji. Background signal was corrected using a rolling ball subtraction algorithm, and a consistent automated threshold (Otsu method) was applied to generate binary masks across all samples. The percent area occupied by GFAP+ signal was calculated as: GFAP % area = (GFAP+ pixels within ROI / total pixels within ROI) × 100. This measurement was performed within each manually delineated ROI. Threshold parameters were held constant within each experimental cohort. Individual GFAP+ astrocyte cell areas (µm²) were quantified to assess cellular hypertrophy at the single-cell level. From the same maximum-intensity projections, individual GFAP+ astrocytes with clearly delineated cell boundaries were identified within each ROI. Cell area was measured by tracing the outer boundary of each GFAP+ cell using the freehand selection tool in Fiji, encompassing the soma and primary process domain (29). Astrocytes with overlapping or ambiguous boundaries were excluded from analysis. Mean cell area was calculated per ROI, averaged across sections, and then per animal for group comparisons.

### Neuron quantification

Neuronal cell density and the thickness of the pyramidal cell layers in CA1 (cornu ammonis 1) and CA2/3 (cornu ammonis 2/3) and the granule cell layer in the dentate gyrus (DG) were quantified in NeuN-immunolabeled sections to assess neuronal cell number and cytoarchitecture across hippocampal subfields. Quantification was performed in Fiji using confocal z-stacks acquired at 20× and converted to maximum-intensity projections. Neuronal density was quantified using a semi-automated dual-channel approach adapted from Woeffler-Maucler et al. (30). This method uses co-labeling of NeuN with the nuclear counterstain DAPI to discriminate NeuN-positive (NeuN⁺) neuronal nuclei from NeuN-negative (NeuN⁻) non-neuronal nuclei within the same field, enabling reliable automated segmentation in densely packed cell layers. The DAPI channel was used to identify all nuclei by automated thresholding and watershed segmentation to separate touching cells. The NeuN channel served as a classification mask: DAPI-positive cells that co-localized with NeuN immunoreactivity above a defined intensity threshold were classified as neuronal (NeuN⁺/DAPI⁺), whereas those below the threshold were classified as non-neuronal (NeuN⁻/DAPI⁺). This dual-channel strategy reduces undercounting that can occur when NeuN-positive cells in tightly packed layers, particularly the hippocampal CA1 and DG, are difficult to resolve by single-channel thresholding alone. [5]. ROI were manually delineated in the principal cell body layers of CA1 (stratum pyramidale), CA2/3 (stratum pyramidale), and DG (stratum granulosum) on each section. ROI areas (mm²) were recorded, and NeuN+ cell counts within each ROI were normalized to area to yield neuronal density, expressed as NeuN+ cells/mm². Thickness of principal cell body layers was measured perpendicular to their long axis in each hippocampal subfield: the pyramidal cell layer (stratum pyramidale) in CA1 and CA2/3, and the granule cell layer (stratum granulosum) in DG. This measure captures thinning of the cell body layer that may result from neuronal loss, dendritic reorganization, or altered packing density, and has been used as an index of neurodegenerative change in hippocampal subfields. For each section, measurements were taken at equidistant points along the cell body layer within each subfield ROI using the straight-line measurement tool in Fiji. The inner and outer boundaries of the layer were identified based on the dense NeuN+ band and the adjacent relatively cell-sparse layers (stratum oriens/stratum radiatum in CA fields; molecular layer/hilus in DG). Measurements were recorded in µm, averaged per subfield per section, and then per animal.

### Behavioral tests

Prior to behavioral assessment, animals were transferred to the testing room and allowed to habituate to environmental conditions for 3-4 hours. All behavior experiments were conducted during the late light phase of the light-dark cycle. After each trial, the apparatus was cleaned with 20% ethanol and thoroughly dried to minimize the residual olfactory cues.

#### Y-maze spontaneous alterations test

Short term spatial working memory was evaluated using a Y-maze consisting of three identical arms (30 × 6 × 15 cm) positioned at 120° angles relative to one another (Panlab Harvard Apparatus, MA, USA) (31, 32). At the beginning of each trial, the mouse was placed at the distal end of one arm facing the center of the maze and allowed to freely explore all three arms for 10 mins. The sequence and total number of arm entries were recorded by experimenters who were blind to genotypes. A single arm entry was defined as the placement of all four limbs within an arm. The percentage of spontaneous alternation [%] was calculated as [Number of consecutive entries into all three arms / (Total number of arm entries − 2)] × 100%.

#### Novel object recognition (NOR) test

NOR test was used to assess object recognition memory and is based on the rodent’s tendency to explore novel objects over familiar objects (31–33). Testing was performed in a white acrylic open-field arena measuring 40 × 40 × 40 cm. The experimental paradigm consisted of three sequential sessions: habituation session, training session and testing session. On the first day, each mouse was individually placed in the empty arena and allowed to freely explore for 5 mins to habituate to the testing environment. On the second day, two identical objects were positioned in the arena. The mouse was placed at the midpoint of the opposite wall and was allowed to freely explore both the objects for 5 mins during the training session. After a 2-hour interval, one of the familiar objects was replaced with a novel object. The mouse was returned to the arena and allowed to explore the objects for 5 mins. The amount of time spent exploring the novel and familiar objects was recorded. The discrimination index was calculated as (novel object exploration time – familiar object exploration time)/ total object exploration time.

### Statistical analysis

Data are presented as mean ± SEM. All statistical analyses were performed using GraphPad Prism 10 (GraphPad Software). The Shapiro-Wilk test was performed to assess the assumption of normality and F test was used to compare variances between groups. For histological measures quantified across multiple brain subfields within the same animals (microglia density, microglia soma area, astrocyte density, and astrocyte cell area), data were analyzed using two-way repeated measures ANOVA followed by Bonferroni’s multiple comparisons test. A two-tailed unpaired t-test was used for comparisons between two groups when the data were normally distributed and had equal variances. The Mann–Whitney U test was used for non-normally distributed data. For within-subject comparisons of exploration time between familiar and novel objects in the novel object recognition test, a two-tailed paired t-test was used. A *P* value < 0.05 was considered statistically significant for all analysis.

## RESULTS

### Regional distribution of DARC/ACKR1 throughout the brain

Defining the anatomical distribution of DARC/ACKR1 is an important first step toward understanding its potential role in regulating chemokine availability and neuroinflammatory processes in the brain. We therefore characterized the regional distribution of DARC/ACKR1 protein across the rostro-caudal extent of the mouse brain and assessed potential sex differences in its expression pattern. Male and female C57BL/6J mice were analyzed by immunofluorescence staining using a specific antibody against DARC/ACKR1. Coronal brain sections spanning the rostro-caudal axis were imaged under identical acquisition conditions, allowing systematic comparison of DARC/ACKR1 immunoreactivity across brain regions and between sexes.

As shown in Figure 1, DARC/ACKR1 immunoreactivity was widely distributed throughout the mouse brain, with substantial regional heterogeneity in expression intensity. The overall distribution pattern and relative intensity of DARC/ACKR1 immunoreactivity were highly similar between male and female mice, with no apparent sex-dependent differences across the rostro-caudal extent of the brain. Across cortical regions, DARC/ACKR1 immunoreactivity was relatively stronger in the orbital and piriform cortices, whereas moderate expression was observed in the prefrontal and anterior cingulate cortices. The hippocampus displayed marked subregional heterogeneity, with the strongest immunoreactivity in CA2, moderate expression in CA3 and DG, and comparatively low expression in CA1 (Figures 1 and 2). Among subcortical structures, high levels of DARC/ACKR1 immunoreactivity were observed in the striatum, thalamus, and substantia nigra, followed by the globus pallidus, lateral and medial septum, and nucleus accumbens (Figures 1, 4, 5 and 6). Moderate immunoreactivity was detected in the amygdala, particularly the medial amygdala, as well as in the superior colliculus and midbrain reticular nucleus, whereas the most subregions of the hypothalamus and ventral tegmental area exhibited relatively low immunoreactivity (Figure 1). In the cerebellum, DARC/ACKR1 immunoreactivity was prominent in the Purkinje cell layer and moderate in the molecular layer (Figures 1 and 7). This broad regional distribution suggests that DARC/ACKR1 may have diverse roles in regulating chemokine availability and neuroimmune signaling across distinct brain circuits.

**Figure 1.**
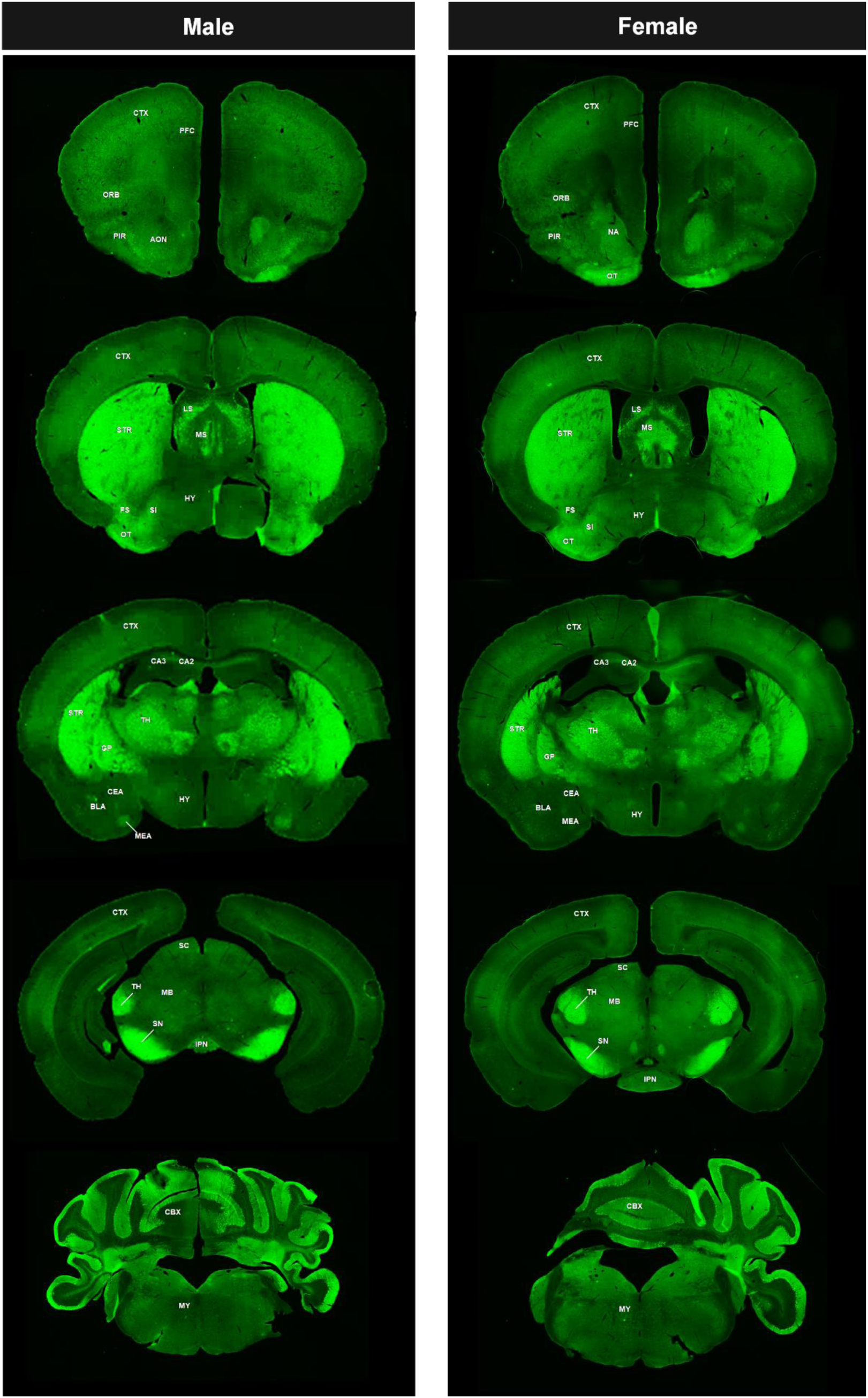
Rostral-to-caudal distribution of DARC/ACKR1 expression in the male and female mouse brain. Representative coronal brain sections spanning the rostral-to-caudal extent of the mouse brain showing the regional distribution and relative abundance of DARC/ACKR1 (green) in male (left) and female (right) mice at 2 months of age. Abbreviations: AN, amygdalar nucleus; AON, anterior olfactory nucleus; BLA, basolateral amygdala; CA2–CA3, Cornu Ammonis field 2–3; CBX, cerebellar cortex; CeA, central amygdala; CTX, cerebral cortex; FS, fundus of striatum; GP, globus pallidus; HY, hypothalamus; IPN, interpeduncular nucleus; LS, lateral septal nucleus; MB, midbrain reticular nucleus; MeA, medial amygdala; MS, medial septal nucleus; MY, medulla; NA, nucleus accumbens; ORB, orbital area; OT, olfactory tubercle; PFC, prefrontal cortex; PIR, piriform cortex; SC, superior colliculus; SI, substantia innominata; SN, substantia nigra; STR, striatum; TH, thalamus.

**Figure 2.**
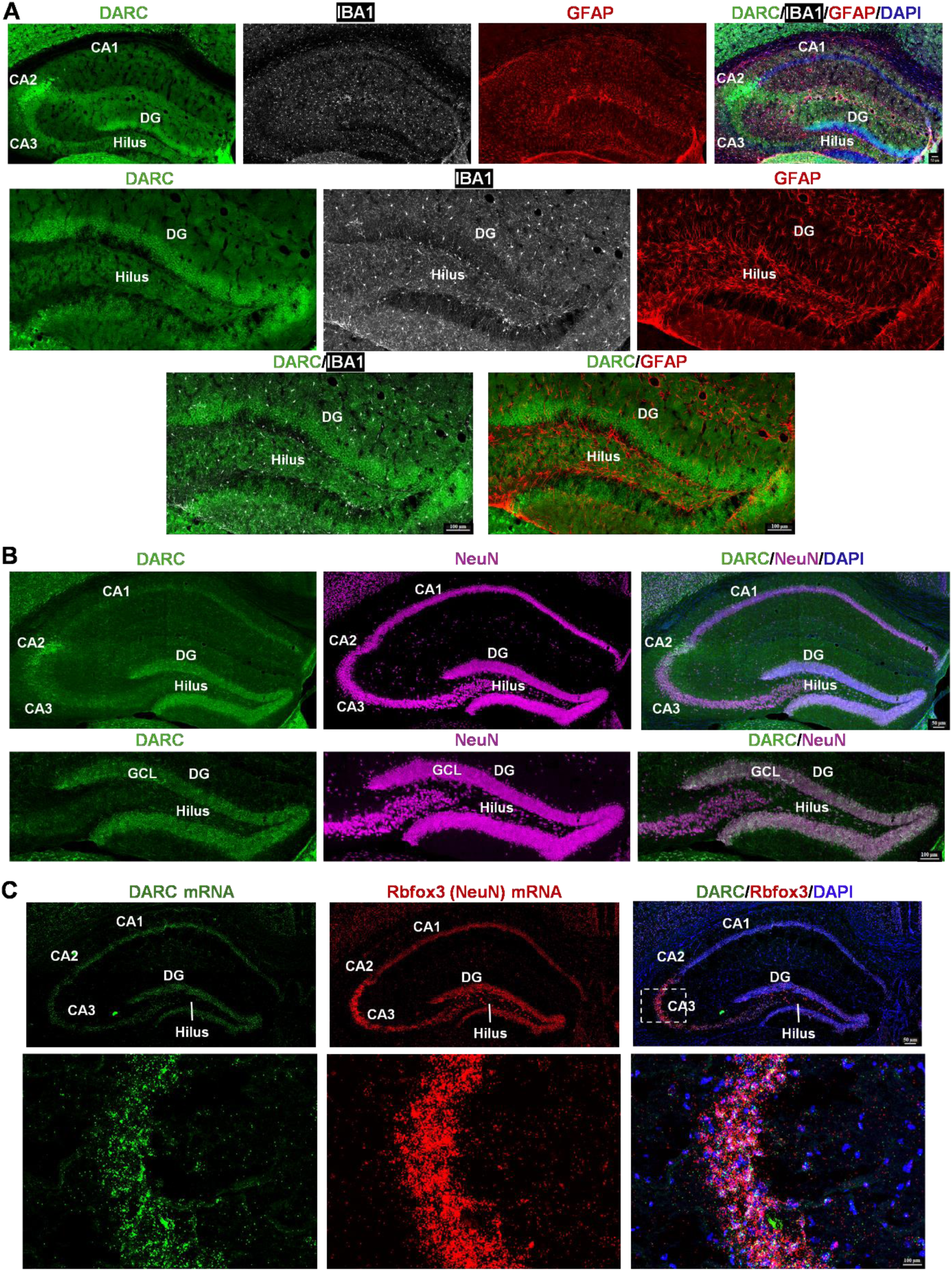
**Cell-type-specific expression of DARC/ACKR1 in the hippocampus**. **A** Confocal images showing triple immunostaining for DARC/ACKR1 (green), the microglial marker IBA1 (white, microglia) and the astrocyte marker GFAP (red, astrocytes) in the hippocampus. Top, images showing DARC/ACKR1, IBA1, and GFAP immunoreactivity across hippocampal subfields CA1–CA3 and the dentate gyrus (DG). Sections were counterstained with DAPI (blue). Middle and bottom, high-magnification images showing DARC/ACKR1, IBA1, and GFAP immunoreactivity in the DG, with no colocalization of DARC/ACKR1 with either IBA1 or GFAP. **B** Confocal images of dual immunostaining for DARC/ACKR1 (green) and the neuronal marker NeuN (magenta, neurons) in the hippocampus. Top, images showing DARC/ACKR1, and NeuN immunoreactivity across CA1–CA3 and DG. Sections were counterstained with DAPI (blue). Bottom, high-magnification images showing co-localization of DARC/ACKR1 and NeuN in the DG. GCL, granule cell layer. **C** RNAscope *in situ* hybridization of mouse brain sections showing expression of *Ackr1* (DARC) mRNA (green) and *Rbfox3* (NeuN) mRNA (red) and their colocalization in the hippocampus. Top, low-magnification images; bottom, high-magnification images of CA3. Sections were counterstained with DAPI (blue).

### Cellular localization of DARC/ACKR1 in the hippocampus and prefrontal cortex

DARC/ACKR1 is an atypical chemokine receptor capable of binding multiple inflammatory chemokines and regulating their availability, distribution, and local concentrations (34–36). Therefore, defining the cellular localization of DARC/ACKR1 in the brain is critical for understanding the cellular context in which it regulates chemokine signaling and neuroinflammatory processes. We first sought to determine whether DARC/ACKR1 is expressed by microglia or astrocytes, the principal resident glial cell populations involved in neuroimmune and neuroinflammatory responses. Because DARC/ACKR1 expression is heterogeneous across the brain, we subsequently examined its cellular localization in functionally distinct brain regions to determine whether its expression pattern varies regionally. In particular, given the proposed role of DARC/ACKR1 in regulating the availability and local distribution of inflammatory CC and CXC chemokines and their signaling through cognate CCR and CXCR receptors (18), we focused on brain regions with substantial expression of these chemokine ligands and receptors and heightened vulnerability to neuroinflammatory insults.

The hippocampus was selected as an initial region of interest because it is highly responsive to chemokine signaling and expresses multiple CC and CXC chemokines and their cognate receptors (37). Previous studies have demonstrated the expression of chemokine receptors, including CCR2, CCR5, CXCR3, and CXCR4, in the adult hippocampus, together with their corresponding chemokine ligands (38–40). We therefore examined the cellular localization of DARC/ACKR1 in the hippocampus. Microglia and astrocytes both produce and respond to inflammatory chemokines and express multiple CCR and CXCR receptors, positioning them to respond to changes in the local chemokine environment (41–43). To determine whether DARC/ACKR1 is expressed by these glial populations, we assessed its colocalization with IBA1, a marker of microglia, and GFAP, a marker of astrocytes. Despite presence of DARC/ACKR1 immunoreactivity across multiple hippocampal subregions, there was no detectable colocalization of DARC/ACKR1 with either IBA1-positive microglia or GFAP-positive astrocytes (Figure 2A). Hippocampal neurons also express multiple chemokine receptors, including members of the CCR and CXCR families, enabling chemokines to directly modulate neuronal signaling and function (44–46). Given the absence of DARC/ACKR1 colocalization with IBA1 or GFAP, we next investigated whether DARC/ACKR1 is expressed by hippocampal neurons. In the DG, DARC/ACKR1 immunoreactivity was distributed throughout the granule cell layer, encompassing both the upper and lower blades, whereas in the CA2 region, immunoreactivity was concentrated within the pyramidal cell layer. Dual immunofluorescence staining with the neuronal marker NeuN demonstrated clear colocalization of DARC/ACKR1 with NeuN-positive neurons (Figure 2B).

To further validate neuronal expression, we performed RNAscope *in situ* hybridization to detect DARC/ACKR1 mRNA together with Rbfox3 mRNA, which encodes the neuronal marker NeuN. Consistent with the immunofluorescence findings, DARC/ACKR1 mRNA signals were detected throughout the CA1–CA3 regions and DG (Figure 2C). DARC/ACKR1 mRNA signals colocalized with *Rbfox3*-positive neurons across these hippocampal subregions (Figure 2C), providing independent evidence that DARC/ACKR1 is expressed by hippocampal neurons.

We next examined the cellular localization of DARC/ACKR1 in the prefrontal cortex (PFC), a brain region that also expresses CC and CXC chemokines and their receptors (47–49). The PFC is a key neuroanatomical substrate for higher-order cognitive processes and is highly vulnerable to neuroinflammatory insults (50–52). Similar to our findings in the hippocampus, DARC/ACKR1 immunoreactivity in the mouse PFC was detected in NeuN-positive cells, with no detectable colocalization with IBA1-positive microglia (Figure 3A). Because GFAP immunoreactivity was extremely low in the PFC of normal mice, astrocytic localization was not further assessed in this region. Consistent with the immunofluorescence findings, RNAscope analysis demonstrated colocalization of DARC/ACKR1 mRNA with Rbfox3 mRNA in the mouse PFC (Figure 3B), further confirming neuronal DARC/ACKR1 expression at the transcript level.

**Figure 3.**
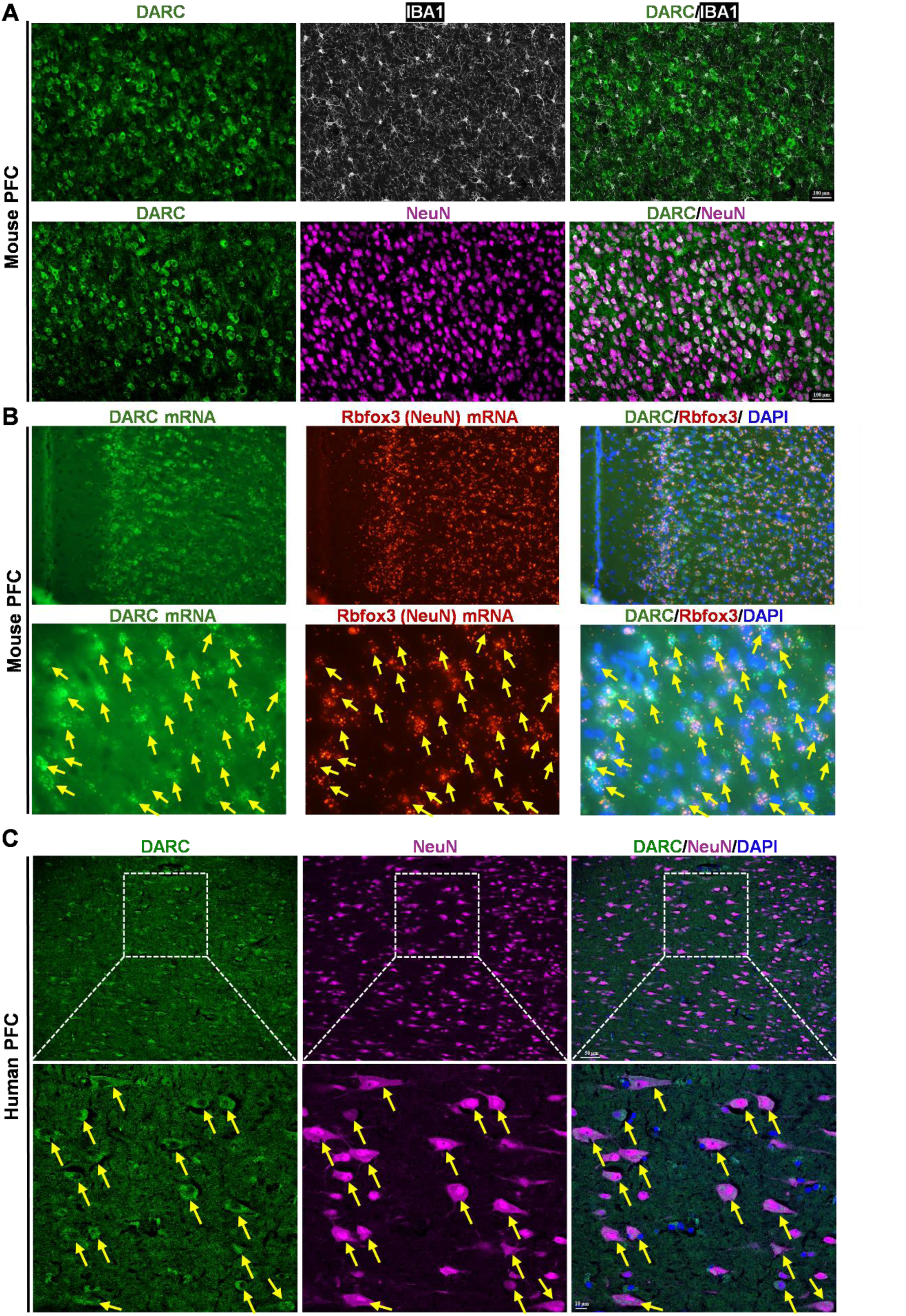
Cell-type-specific expression of DARC/ACKR1 in the mouse and human prefrontal cortex (PFC). **A** Mouse PFC. Top, confocal images showing co-immunostaining for DARC/ACKR1 (green) and the microglial marker IBA1 (white). Bottom, confocal images showing dual immunostaining for DARC/ACKR1 (green) and the neuronal marker NeuN (magenta, neurons) and their colocalization. **B** RNAscope *in situ* hybridization of mouse brain sections showing expression of *Ackr1* (DARC) mRNA (green) and *Rbfox3* (NeuN) mRNA (red) and their colocalization in the PFC. Top, low-magnification images; bottom, high-magnification images. Sections were counterstained with DAPI (blue). **C** Human PFC. Representative confocal images of human brain tissue from dorsolateral PFC (Brodmann area 9, BA9) showing dual immunostaining of DARC/ACKR1 (green) and NeuN (magenta, neurons), with merged images illustrating their colocalization.

We next examined whether the neuronal expression pattern of DARC/ACKR1 observed in the mouse brain is conserved in humans. Human dorsolateral PFC (DLPFC; Brodmann area 9) samples from six neurologically normal subjects were analyzed by dual immunofluorescence staining for DARC/ACKR1 and NeuN. DARC/ACKR1 immunoreactivity was localized to NeuN-positive neurons throughout the examined region. Notably, nearly all DARC/ACKR1-positive cells were NeuN-positive, indicating a neuronal predominance of DARC/ACKR1 expression in the human DLPFC (Figure 3C).

### Cellular localization of DARC/ACKR1 in the septum, striatum, thalamus, and cerebellum

We next examined additional brain regions with relatively high DARC/ACKR1 expression, including the lateral septum, striatum, thalamus, and cerebellum. Because these regions differ substantially in cellular composition, anatomical organization, and functional properties, we examined DARC/ACKR1 localization across these anatomically and functionally distinct regions to determine whether its predominant neuronal expression represents a general feature of DARC/ACKR1 distribution throughout the brain rather than a region-specific pattern. In the lateral septum, DARC/ACKR1 immunoreactivity showed no detectable colocalization with IBA1-positive microglia or GFAP-positive astrocytes (Figure 4A). DARC/ACKR1-positive cells exhibited neuronal morphology, and the majority were NeuN-positive (Figure 4B). A subset of DARC/ACKR1-positive cells, however, lacked detectable NeuN immunoreactivity. Nevertheless, the absence of DARC/ACKR1 colocalization with microglial or astrocytic markers, together with the predominance of NeuN-positive DARC/ACKR1-positive cells, supports predominantly neuronal localization of DARC/ACKR1 in the lateral septum. A similar pattern was observed in the striatum and thalamus. In the striatum, the vast majority of DARC/ACKR1-positive cells were NeuN-positive (Figure 5B), with no detectable colocalization with IBA1-positive microglia or GFAP-positive astrocytes (Figure 5A). Likewise, in the thalamus, DARC/ACKR1-positive cells were predominantly NeuN-positive (Figure 6B) and showed no apparent colocalization with either IBA1 or GFAP (Figure 6A). In the cerebellum, strong DARC/ACKR1 immunoreactivity was observed within the Purkinje cell layer, DARC/ACKR1-positive cells exhibited characteristic Purkinje cell morphology (Figure 7A). Interestingly, DARC/ACKR1-positive Purkinje cells were NeuN-negative but expressed GAD67 (Figure 7B), consistent with their GABAergic neuronal identity. This finding suggests that NeuN is not uniformly detectable across neuronal populations and that the absence of NeuN immunoreactivity should not, by itself, be interpreted as evidence of a non-neuronal identity. Collectively, our findings across multiple brain regions demonstrate that DARC/ACKR1 is exclusively expressed by neurons, with no detectable expression in microglia or astrocytes in the brain regions examined.

**Figure 4.**
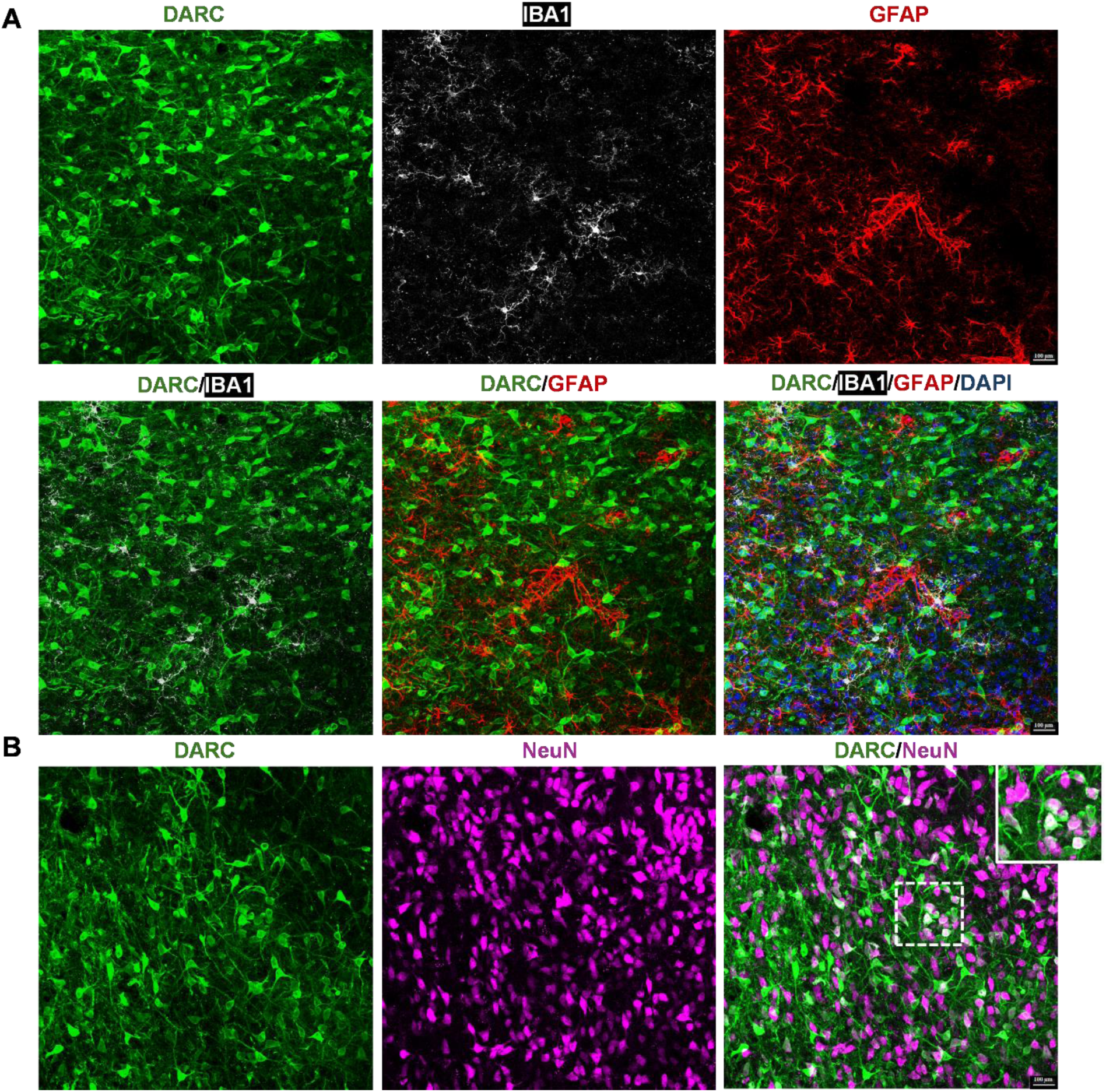
Cell-type-specific expression of DARC/ACKR1 in the lateral septum. **A** Top, confocal images showing triple immunostaining for DARC/ACKR1 (green), the microglial marker IBA1 (white), and the astrocyte marker GFAP (red) in the lateral septum. Individual channels are shown separately. Bottom, merged images showing no colocalization of DARC/ACKR1 with either IBA1 or GFAP. **B** Confocal images showing dual immunostaining for DARC/ACKR1 (green) and the neuronal marker NeuN (red) in the lateral septum. Merged images showing colocalization of DARC/ACKR1 with NeuN in lateral septal neurons. Inset, higher-magnification view illustrating DARC/ACKR1 and NeuN colocalization.

**Figure 5.**
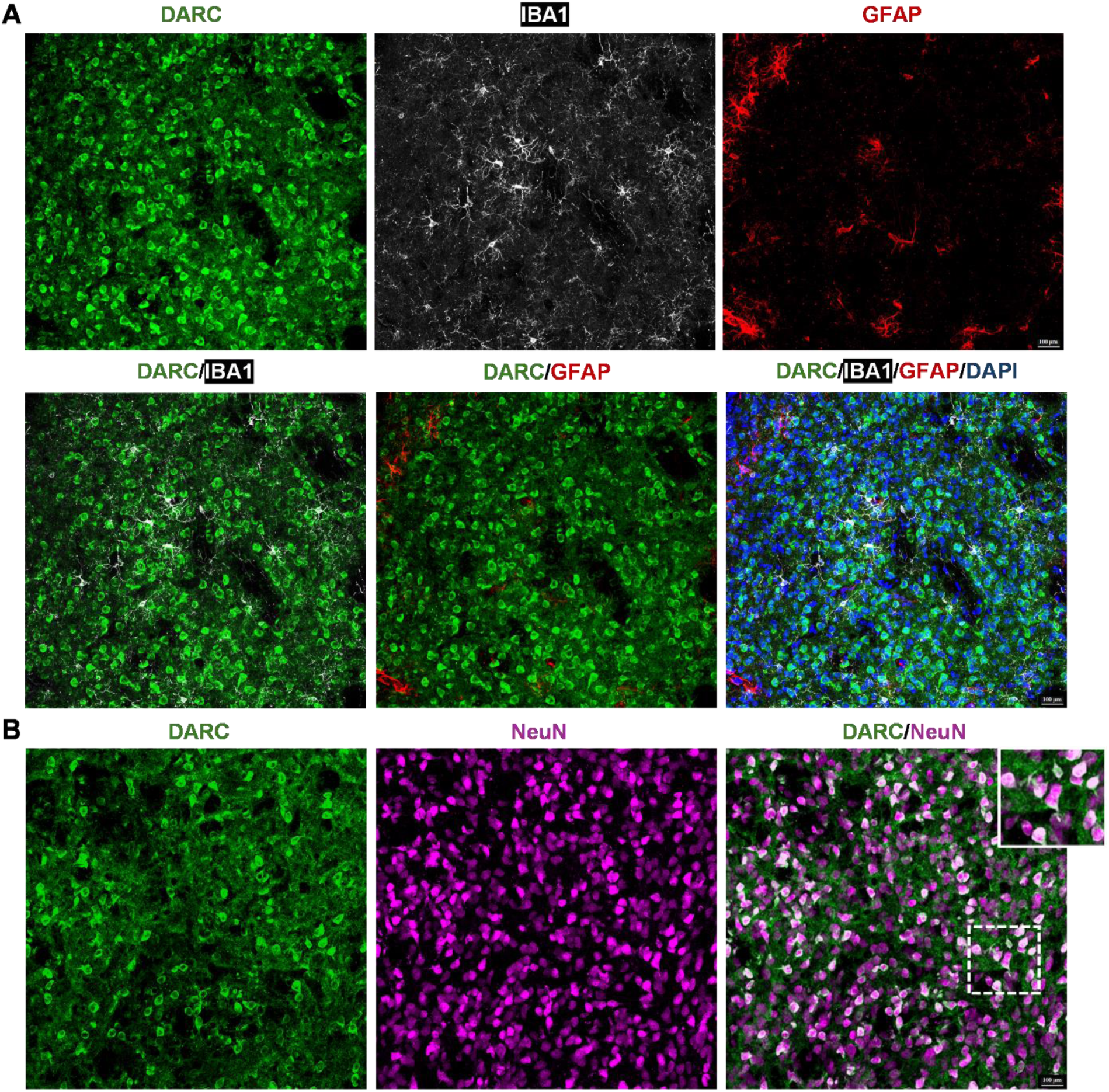
Cell-type-specific expression of DARC/ACKR1 in the striatum. **A** Top, confocal images showing triple immunostaining for DARC/ACKR1 (green), the microglial marker IBA1 (white), and the astrocyte marker GFAP (red) in the striatum. Individual channels are shown separately. Bottom, merged images showing no colocalization of DARC/ACKR1 with either IBA1 or GFAP. **B** Confocal images showing dual immunostaining for DARC/ACKR1 (green) and the neuronal marker NeuN (red) in the striatum. Merged images showing colocalization of DARC/ACKR1 with NeuN in striatal neurons. Inset, higher-magnification view illustrating DARC/ACKR1 and NeuN colocalization.

**Figure 6.**
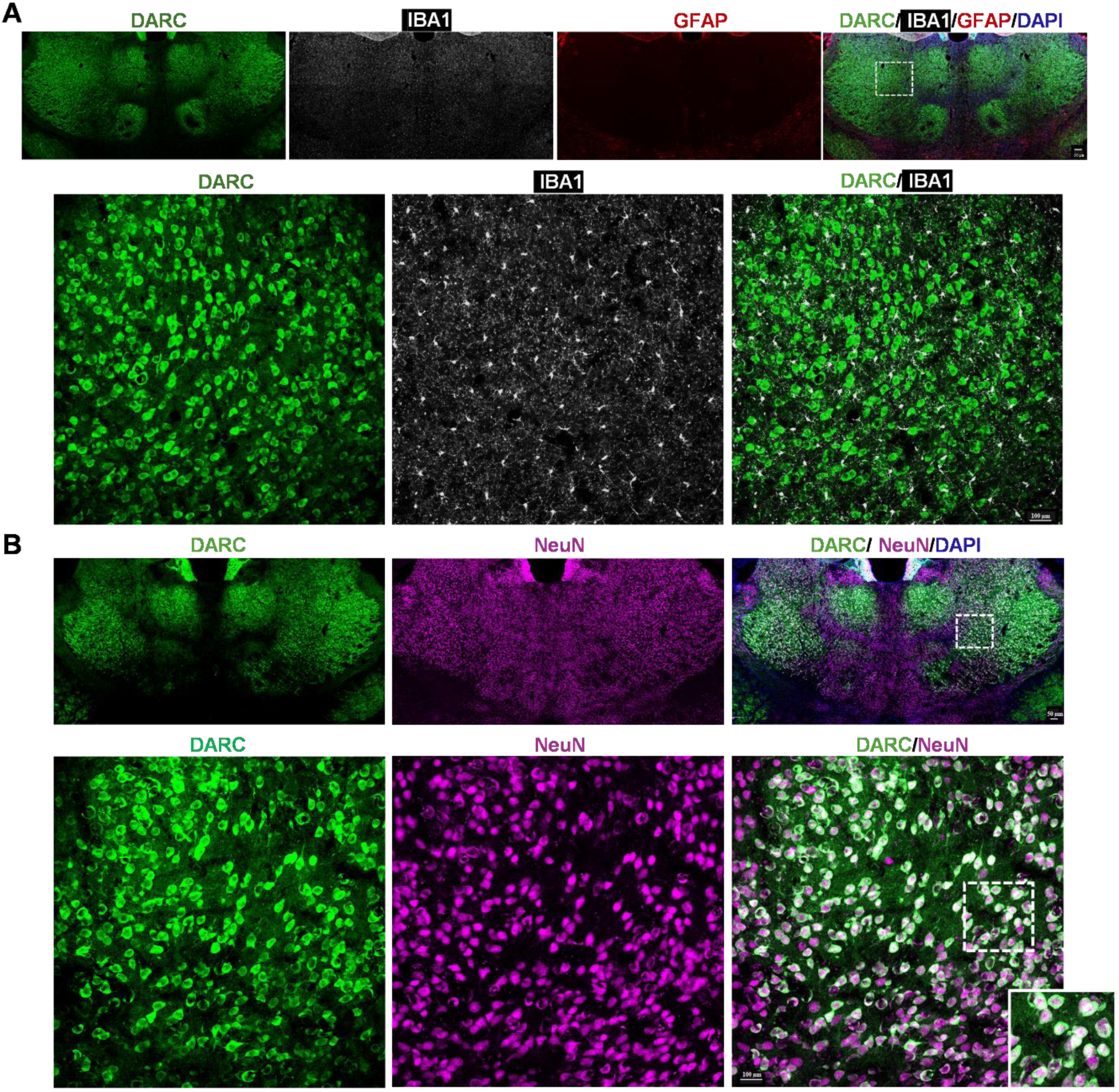
Cell-type-specific expression of DARC/ACKR1 in the thalamus. **A** Top, confocal images showing triple immunostaining for DARC/ACKR1 (green), the microglial marker IBA1 (white), and the astrocyte marker GFAP (red) in the thalamus. Individual channels are shown separately. Bottom, merged images showing no colocalization of DARC/ACKR1 with IBA1. **B** Confocal images showing dual immunostaining for DARC/ACKR1 (green) and the neuronal marker NeuN (red) in the thalamus. Top, low-magnification images showing DARC/ACKR1 and NeuN immunostaining; bottom, high-magnification images showing DARC/ACKR1 and NeuN colocalization in thalamic neurons. Inset, higher-magnification view illustrating DARC/ACKR1 and NeuN colocalization.

### DARC/ACKR1 deficiency leads to increased microglial activation and astrogliosis in the brain

To determine whether DARC/ACKR1 is critical for controlling the neuroinflammatory state, we examined microglial and astrocytic changes in the hippocampus of DARC/ACKR1-deficient mice and their WT littermates. Compared with WT littermate mice, which exhibited low levels of microglial and GFAP immunoreactivity, DARC/ACKR1-deficient mice showed marked increases in microglial and astrocytic responses across hippocampal subregions (Figure 8A and B). Quantitative analysis revealed a significant increase in microglial density in the CA1, CA2/3, and DG regions of DARC/ACKR1-deficient mice (Figure 8C). In addition, microglial soma area was significantly increased across these hippocampal regions (Figure 8C). Increased number and enlarged soma size of microglia indicate robust microglial activation in the hippocampus in mice with genetic deletion of DARC/ACKR1. Similarly, DARC/ACKR1-deficient mice exhibited pronounced astrogliosis. The density of GFAP-positive astrocytes was significantly increased in the CA1, CA2/3, and DG regions, accompanied by an increase in astrocyte cell area across these subregions (Figure 8). The concurrent increase in GFAP-positive cell density and cellular area is indicative of marked astrocytic activation and hypertrophy. Together with the observed increases in microglial density and soma area, these findings demonstrate that genetic deletion of DARC/ACKR1 is associated with widespread alterations in microglial and astrocytic responses across the hippocampus. We also examined the expression of CCL2/MCP-1, a ligand of DARC/ACKR1, in the hippocampus. As a preliminary observation, CCL2/MCP-1 expression appeared to be markedly increased throughout the hippocampus of DARC/ACKR1-deficient mice compared with WT mice (Supplementary Figure 1). This increase in CCL2/MCP-1 may further contribute to the neuroinflammatory consequences of DARC/ACKR1 deficiency.

**Figure 7.**
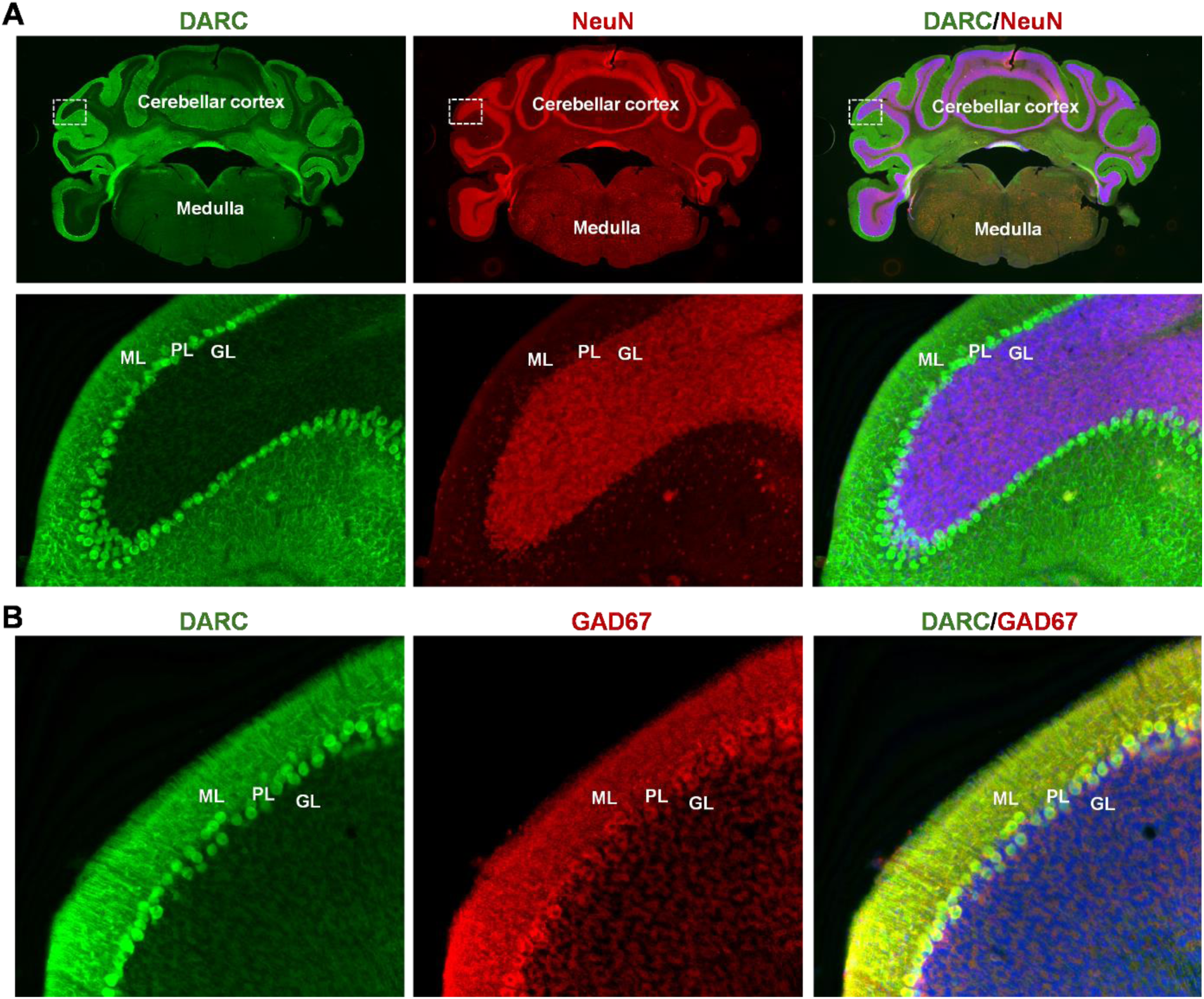
Expression of DARC/ACKR1 by GABAergic Purkinje cells in the cerebellum. **A** Confocal image showing dual immunostaining for DARC/ACKR1 (green) and the neuronal marker NeuN (red) in the cerebellar cortex. Top, low-magnification images showing DARC/ACKR1 and NeuN immunoreactivity; bottom, higher-magnification images showing that NeuN does not label Purkinje cells. **B** Confocal images showing dual immunostaining for DARC/ACKR1 (green) and the GABAergic neuronal marker GAD67 (red) in the cerebellum. Merged images show colocalization of DARC/ACKR1 and GAD67 in Purkinje cells. GL, granular layer; ML, molecular layer; PL, Purkinje cell layer.

**Figure 8.**
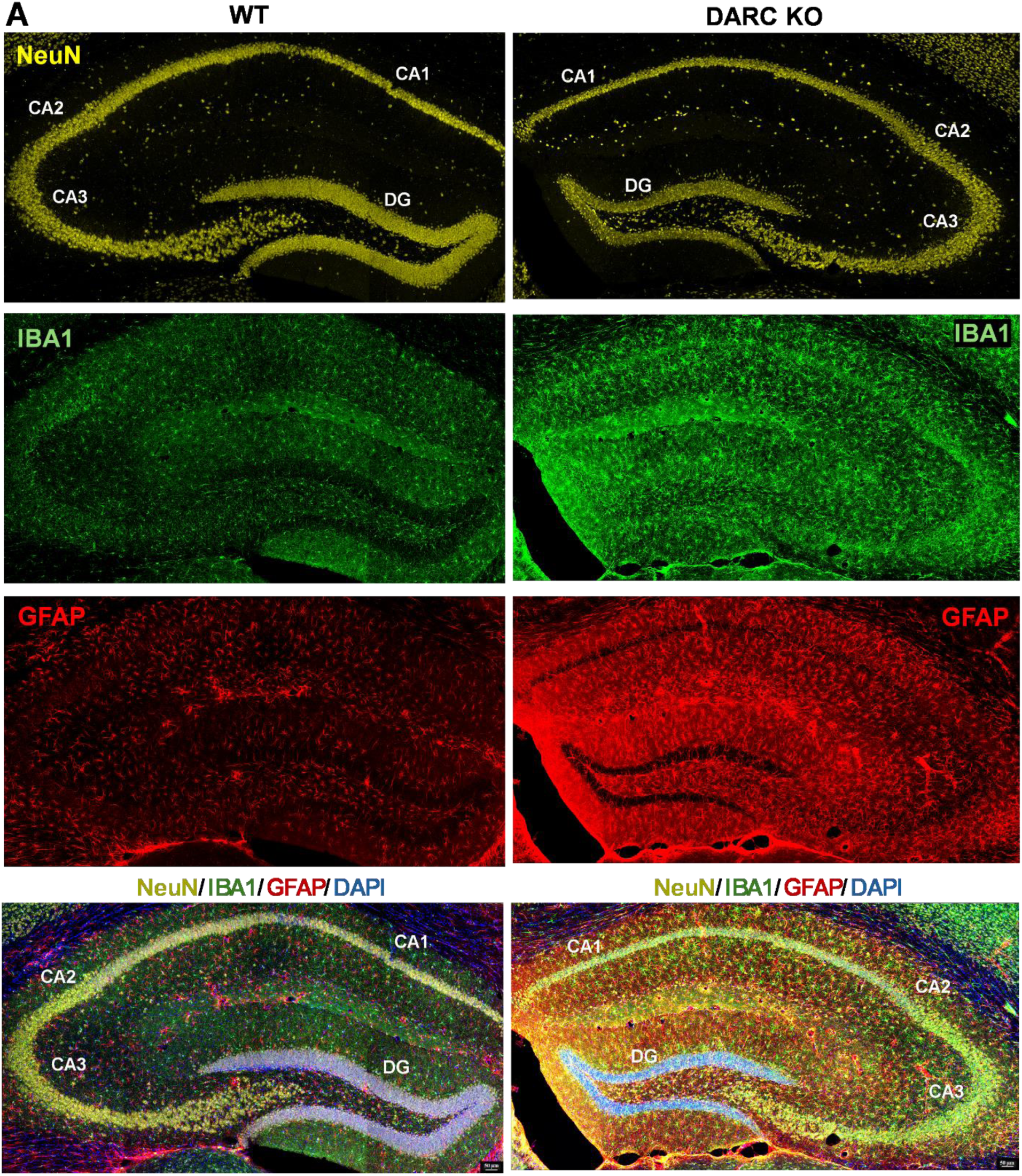
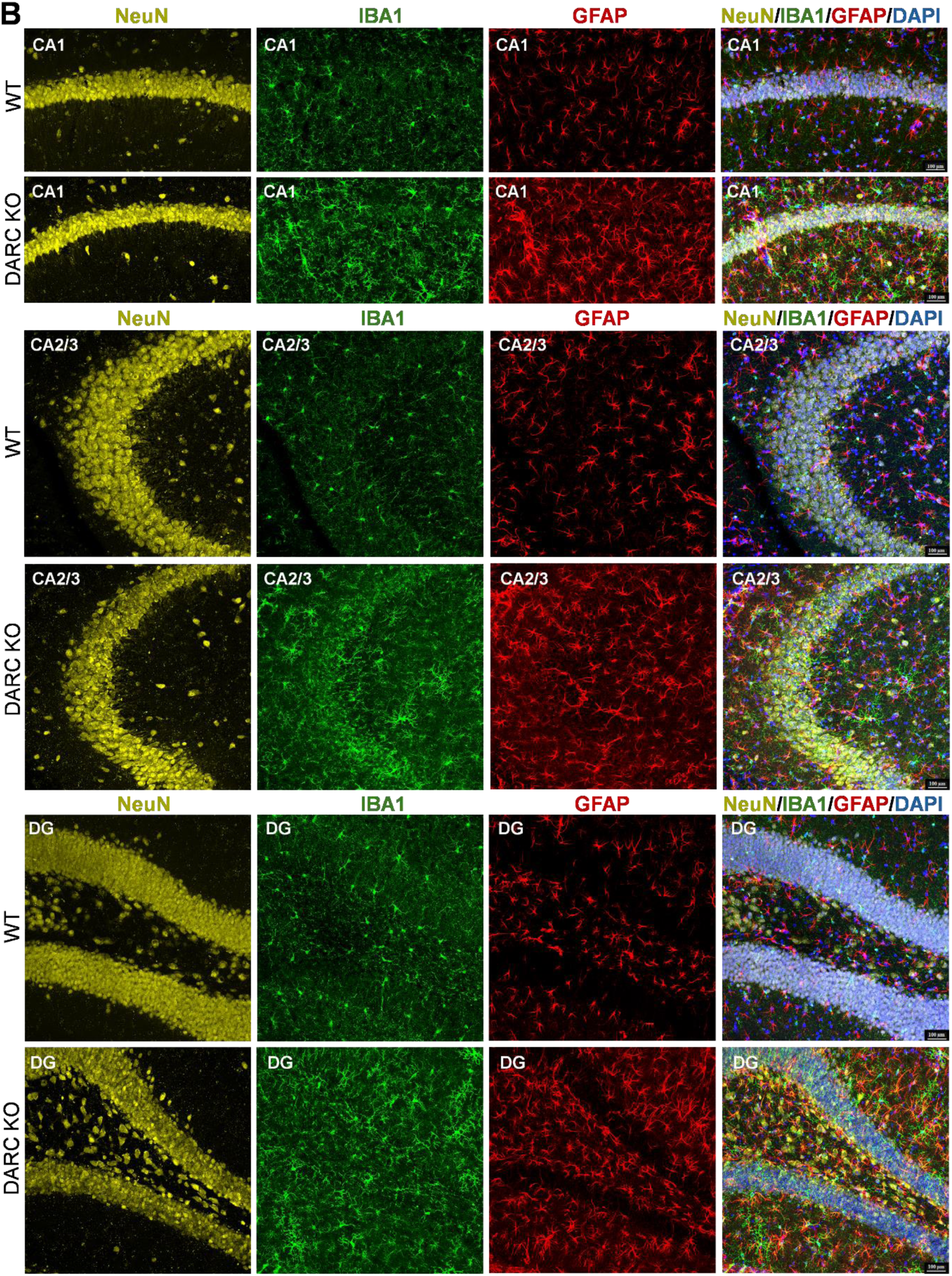
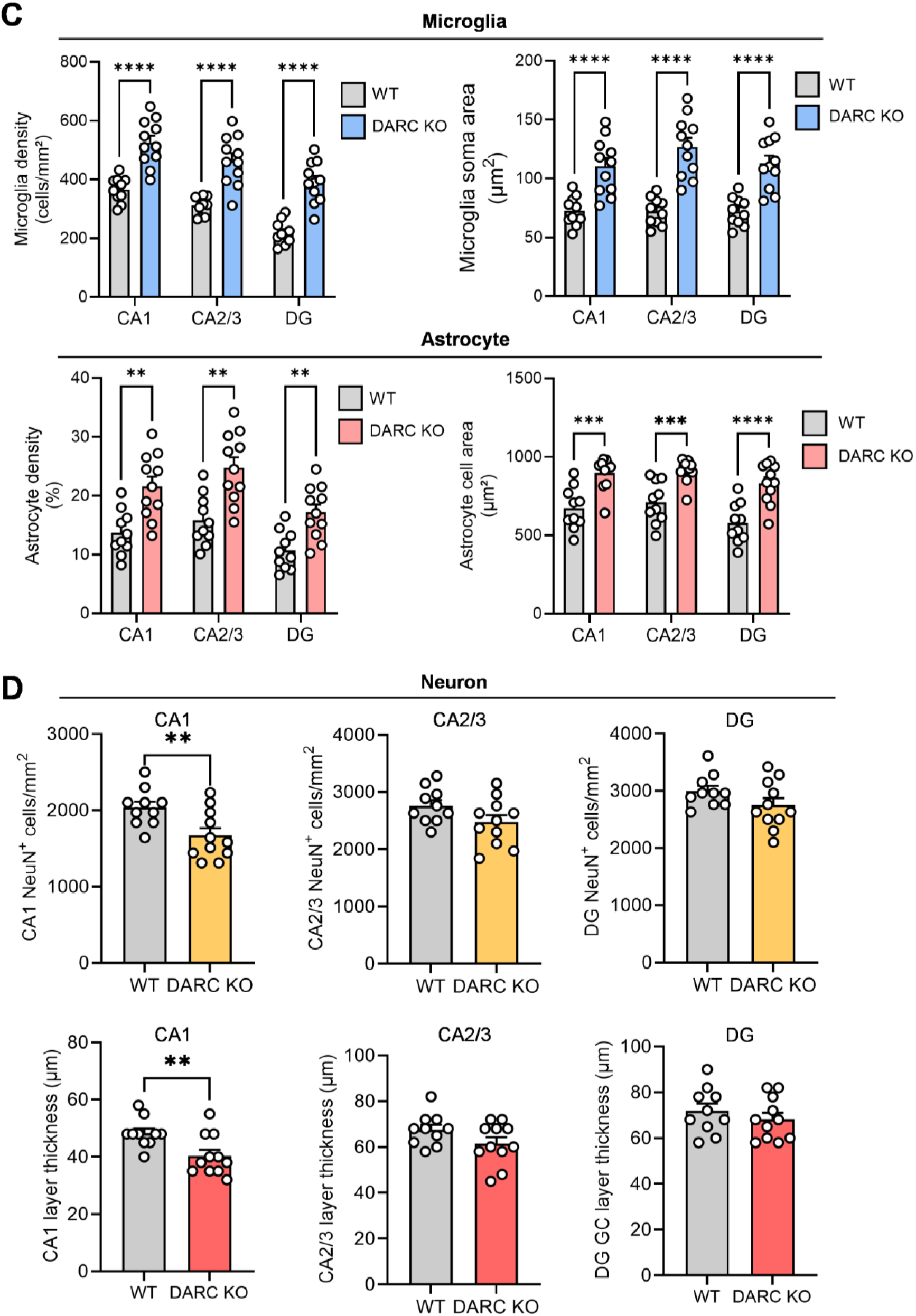
DARC/ACKR1 deficiency induces neuroinflammatory responses and neuronal degeneration across hippocampal subfields in DARC/ACKR1-deficient mice. **A** Representative low-magnification confocal images showing triple immunostaining for NeuN (yellow, neurons), IBA1 (green, microglia), and GFAP (red, astrocytes), with DAPI counterstaining, in the hippocampus of wild-type (WT) and DARC/ACKR1-deficient (DARC KO) mice. Individual channels are shown separately, and merged images with DAPI counterstaining (blue) are shown in the bottom row. **B** High-magnification confocal images showing triple immunostaining for NeuN, IBA1, and GFAP in the CA1, CA2/3, and dentate gyrus (DG) subfields of the hippocampus in WT and DARC/ACKR1 KO mice. **C** Quantification of microglial, astrocytic, and neuronal changes across hippocampal subfields in WT and DARC/ACKR1 KO mice. Top, microglial density (cells/mm²) and soma area (µm²). For microglial density, two-way ANOVA revealed significant effects of genotype (*F*(_1,19_) = 36.27, *P* < 0.0001) and hippocampal subregion (*F*_(2,38)_ = 568.3, *P* < 0.0001), with no significant genotype × subregion interaction (*F*_(2,38)_ = 2.352, *P* = 0.1089). Post hoc comparisons showed significantly higher microglial density in DARC/ACKR1 KO mice than in WT mice in CA1, CA2/3, and DG (all *P* < 0.0001). For microglial soma area, two-way ANOVA revealed significant effects of genotype (*F*_(1,19)_ = 29.14, *P* < 0.0001) and hippocampal subregion (*F*_(2,38)_ = 22.12, *P* < 0.0001), as well as a significant genotype × subregion interaction (*F*_(2,38)_ = 23.73, *P* < 0.0001). Post hoc comparisons showed significantly larger microglial soma area in DARC/ACKR1 KO mice than in WT mice in CA1, CA2/3, and DG (all *P* < 0.0001). Bottom, astrocyte density (%) and cell area (µm²). For astrocyte density, two-way ANOVA revealed significant effects of genotype (*F*_(1,19)_ = 14.65, *P* = 0.0011) and hippocampal subregion (*F*_(2,38)_ = 451.9, *P* < 0.0001), as well as a significant genotype × subregion interaction (*F*_(2,38)_ = 16.39, *P* < 0.0001). Post hoc comparisons showed significantly higher astrocyte density in DARC/ACKR1 KO mice than in WT mice in CA1 (*P* = 0.0034), CA2/3 (*P* = 0.0027), and DG (*P* = 0.0038). For astrocyte cell area, two-way ANOVA revealed significant effects of genotype (*F*_(1,19)_ = 21.18, *P* = 0.0002) and hippocampal subregion (*F*_(2,38)_ = 47.04, *P* < 0.0001), but no significant genotype × subregion interaction (*F*_(2,38)_ = 2.636, *P* = 0.0847). Post hoc comparisons showed significantly larger astrocyte cell area in DARC/ACKR1 KO mice than in WT mice in CA1 (*P* = 0.0001), CA2/3 (*P* = 0.0007), and DG (*P* < 0.0001). **D** Quantification of neuronal changes across hippocampal subfields in WT and DARC/ACKR1 KO mice. Top, neuronal cell density (cells/mm²). Independent *t*-tests showed a significant difference between WT and DARC/ACKR1 KO mice in CA1 (*t*₁₉ = 2.927, *P* = 0.0087), but not in CA2/3 (*t*₁₉ = 1.799, *P* = 0.0879) or DG (*t*₁₉ = 1.578, *P* = 0.1311). Bottom, neuronal layer thickness (µm). A significant difference was detected in CA1 (*t*₁₉ = 2.933, *P* = 0.0085), but not in CA2/3 (*t*₁₉ = 1.670, *P* = 0.1113) or DG (*t*₁₉ = 0.8623, *P* = 0.3933). WT: n = 10 mice; DARC KO: n = 11 mice. \**P* < 0.05, \*\**P* < 0.01, \*\*\**P* < 0.001, \*\*\*\**P* < 0.0001 compared with the WT group. Data are presented as mean ± SEM.

Microglial activation and astrogliosis were also evaluated in the lateral septum and striatum to determine whether the neuroinflammatory phenotype associated with ACKR1 deficiency extended to additional brain regions. In the lateral septum, DARC/ACKR1-deficient mice exhibited clear evidence of microglial and astrocytic activation. Microglial density and soma area were significantly increased, indicating both an expansion of the microglial population and somatic enlargement (Figure 9A). Similarly, GFAP staining revealed significant increases in both astrocyte density and cell area, indicating an increased astrocyte population together with cellular hypertrophy, consistent with enhanced astrogliosis in this region (Figure 9A). In the striatum, DARC/ACKR1 deficiency produced similar changes in both microglia and astrocytes. Microglial density and soma area were significantly increased (Figure 9B), consistent with microglial activation, while GFAP-positive astrocytes showed significant increases in both density and cell area, indicating astrocyte expansion and hypertrophy (Figure 9B). Collectively, these findings demonstrate that DARC/ACKR1 deficiency induces microglial activation and astrogliosis beyond the hippocampus, with the cellular features and magnitude of these responses varying across brain regions.

**Figure 9.**
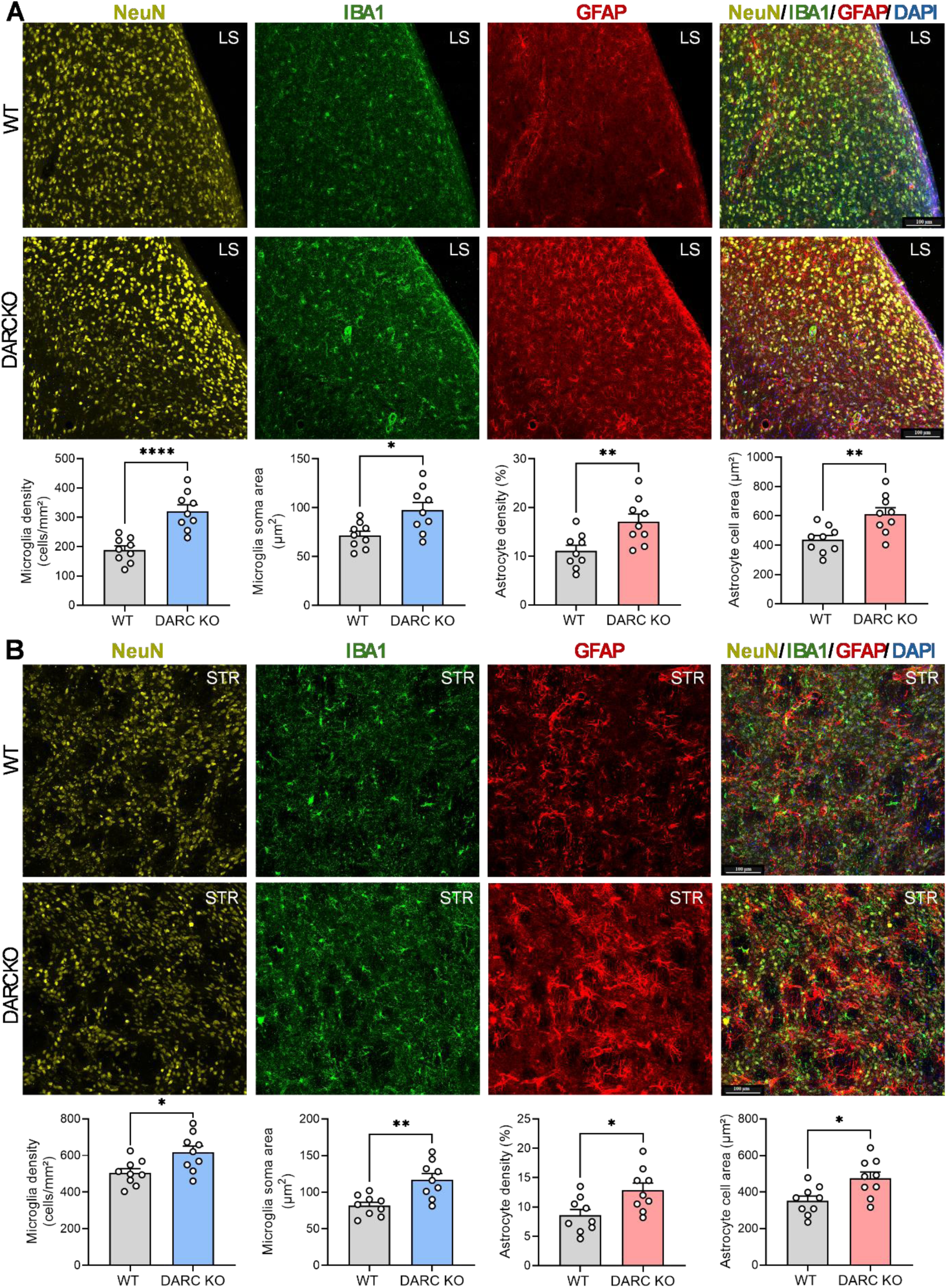
DARC/ACKR1 deficiency induces neuroinflammatory responses in the lateral septum and striatum. Microglial, astrocytic, and neuronal changes were assessed in the lateral septum and striatum of DARC/ACKR1-deficient (DARC KO) mice and their WT littermate controls. **A** Representative low-magnification confocal images showing triple immunostaining for NeuN (yellow, neurons), IBA1 (green, microglia), and GFAP (red, astrocytes), with DAPI counterstaining, in the lateral septum (LS) of WT (top row) and DARC KO (middle row) mice. Bottom, Quantification of microglial and astrocytic changes in the DARC KO mice compared with WT littermates. Left, microglia density (cells/mm²; *t*₁₆ = 5.179, *P* < 0.0001); middle-left, microglial soma area (µm²; *t*₁₆ = 2.915, *P* = 0.0101); middle-right, astrocyte density (cells/mm²; *t*₁₆ = 3.092, *P* = 0.0070); right, astrocyte cell area (µm²; *t*₁₆ = 3.268, *P* = 0.0048). **B** Representative low-magnification confocal images showing triple immunostaining for NeuN (yellow, neurons), IBA1 (green, microglia), and GFAP (red, astrocytes), with DAPI counterstaining, in the striatum (Str) of WT (top row) and DARC KO (middle row) mice. Bottom, Quantification of microglial and astrocytic changes in the DARC KO mice compared with WT littermates. Left, microglia density (cells/mm²; *t*_16_ = 2.703, *P* = 0.0157); middle-left, microglial soma area (µm²; *t*_16_ = 3.738, *P* = 0.0018); middle-right, astrocyte density (cells/mm²; *t*_16_ = 2.744, *P* = 0.0144); right, astrocyte cell area (µm²; *t*_16_ = 2.867, *P* = 0.0112). WT: n = 9 mice. DARC KO: n = 9 mice. *P < 0.05, **P < 0.01, ****P < 0.0001 compared with the WT group. Data are presented as mean ± SEM.

### DARC/ACKR1 deficiency results in alterations in hippocampal neuronal density and cytoarchitecture

Given the neuronal localization of DARC/ACKR1, we next examined whether DARC/ACKR1 deficiency is associated with alterations in hippocampal neuronal density and neuronal organization. In the CA1 region, DARC/ACKR1 deficiency significantly reduced NeuN-positive neuronal cell density and concomitantly decreased the thickness of the CA1 pyramidal cell layer (Figure 8D, left panel), indicating alterations in the neuronal population and cytoarchitecture of this region. In contrast, neither NeuN-positive neuronal density nor pyramidal layer thickness was significantly altered in the CA2/CA3 regions (Figure 8D, middle panel). Similarly, in the DG, the number of NeuN-positive neurons and the thickness of the granule cell layer were comparable between genotypes (Figure 8D, right panel). Together, these findings suggest that hippocampal subregions exhibit differential vulnerability to DARC/ACKR1 deficiency, with CA1 being particularly susceptible to alterations in neuronal density and cytoarchitecture.

### DARC/ACKR1 deficiency impairs cognitive function

To determine whether genetic deletion of DARC/ACKR1 affects cognitive function, DARC/ACKR1-deficient and WT littermate mice were assessed using the Y-maze and novel object recognition (NOR) tests. In the Y-maze, DARC/ACKR1-deficient mice exhibited a significant reduction in spontaneous alternation compared with WT mice (Figure 10A), indicating impaired spatial working memory. Total arm entries did not differ between genotypes (Figure 10A), suggesting that the reduced alternation performance was not attributable to differences in locomotor activity or general exploratory behavior. Cognitive function was further assessed using the NOR test. During the test phase conducted 2 h after training, WT mice showed a significant preference for the novel object over the familiar object, whereas DARC/ACKR1-deficient mice failed to exhibit a significant preference for the novel object (Figure 10B left), indicating impaired recognition memory. Total object investigation time was comparable between genotypes, suggesting that the deficit in novel object preference was not due to reduced object exploration (Figure 10B middle). Consistent with these findings, the discrimination index was significantly lower in DARC/ACKR1-deficient mice than in WT littermates (Figure 10B right). Together, these findings demonstrate that genetic deletion of DARC/ACKR1 is associated with impairments in both spatial working memory and recognition memory.

**Figure 10.**
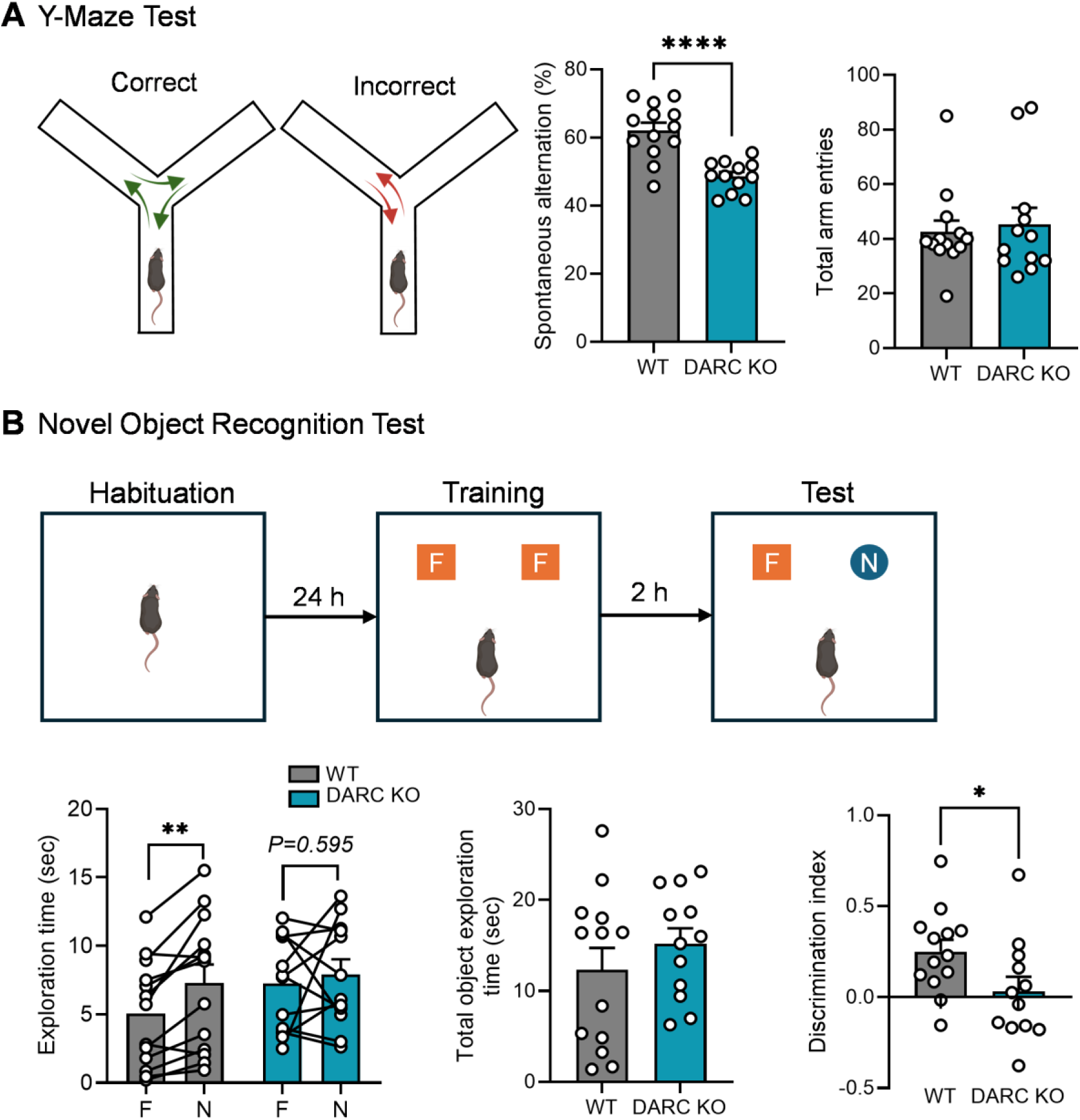
DARC/ACKR1 deficiency impairs spatial working memory and recognition memory. Adult DARC/ACKR1-deficient (DARC KO) mice and their wild-type (WT) littermates, aged 2–8 months, were subjected to two cognitive behavioral tests. **A Y-maze spontaneous alternation test.** Left, schematic illustrating spontaneous alternation, defined as sequential entry into three different arms without re-entering a previously visited arm. Middle, percentage of spontaneous alternation. *t*₂₃ = 5.141, *P* < 0.0001. Right, total arm entries. Total arm entry: Mann– Whitney U test, *P* = 0.8203. Wild-type group (WT) group: n = 13 mice. DARC/ACKR1 deficient (DARC KO) group: n = 12 mice. \*\**P* < 0.001 compared with WT littermate controls. **B Novel object recognition test.** Top, schematic of the experimental timeline: habituation (24 h), training with two identical familiar objects (F), followed by a test session 2 h after training, during which one familiar (F) and one novel (N) object were presented. Bottom, quantification of object investigation. Left, exploration time of familiar versus novel objects (WT, *t*_12_ = 3.992, *P* = 0.0018; DARC KO, *t*_11_ = 0.5468, *P* = 0.5954). Middle, total object exploration time for novel and familiar objects (*t*_23_ = 0.9504, *P* = 0.3518). Right, discrimination index (*t*_23_ = 2.137, *P* = 0.0435). WT, *n* = 13; DARC KO, *n* = 12. \**P* < 0.05, \*\**P* < 0.01, and \*\*\*\**P* < 0.0001 versus the WT group. Data are presented as mean ± SEM. F, familiar object; N, novel object.

## DISCUSSION

Our finding that DARC/ACKR1 is selectively expressed by subsets of neurons, but not microglia or astrocytes, throughout the brain identifies neurons as a previously unrecognized cellular population through which DARC/ACKR1 may regulate local chemokine signaling in the brain. This finding is particularly intriguing given that chemokines have functions beyond their canonical roles in neuroimmune and inflammatory responses, including the regulation of neuronal activity, synaptic transmission and plasticity, and neuron–glia communication (53, 54). As an atypical chemokine receptor capable of binding and sequestering inflammatory chemokines, neuronal DARC/ACKR1 may regulate the local availability and distribution of chemokines, thereby fine-tuning chemokine signaling in neurons and neighboring glial cells. Such regulation could contribute to the maintenance of neuroimmune homeostasis by limiting excessive or sustained chemokine signaling. Importantly, the demonstration of neuronal DARC/ACKR1 expression in the human brain further supports the potential translational relevance of this neuronal chemokine regulatory mechanism.

Our analysis of the regional and cellular distribution of DARC/ACKR1 throughout the brain revealed distinct patterns of neuronal expression. Within the hippocampus, DARC/ACKR1 expression was most prominent in CA2, followed by the DG and CA3, whereas relatively low expression was detected in CA1. DARC/ACKR1 was also abundantly expressed in the lateral septum, striatum, thalamus, and cerebellum. However, the regional abundance of DARC/ACKR1 did not appear to correspond directly to the magnitude of the neuroinflammatory response following its deletion. Despite substantial differences in basal DARC/ACKR1 expression among hippocampal subregions, DARC/ACKR1-deficient mice exhibited robust neuroinflammatory responses throughout CA1–CA3 and the DG, with increased microglial density and soma size and marked astrogliosis across all hippocampal subregions, including CA1, where basal DARC/ACKR1 expression was relatively low. Similarly, brain regions with abundant DARC/ACKR1 expression, such as the lateral septum and striatum, did not exhibit proportionally greater inflammatory responses following DARC/ACKR1 deletion. These findings indicate that the neuroinflammatory consequences of DARC/ACKR1 deficiency are not simply determined by its basal abundance within a given brain region, but may instead reflect regional differences in the susceptibility to disruption of chemokine homeostasis. Indeed, the hippocampus is particularly sensitive to a variety of physiological and pathological insults that can promote microglial activation and astrogliosis (37, 55). One potential mechanism is that neuronal DARC/ACKR1 regulates the extracellular chemokine milieu in a non-cell-autonomous manner. Loss of DARC/ACKR1-mediated chemokine sequestration could increase the local availability of inflammatory chemokines, thereby enhancing chemokine signaling beyond DARC/ACKR1-expressing neurons and promoting or amplifying microglial and astrocytic activation in neighboring cells. The magnitude of this response may also depend on the expression of specific DARC/ACKR1-binding chemokines and their cognate classical chemokine receptors, as individual chemokines differ in their capacity to promote neuroinflammation (56–58). MCP-1/CCL2 is a well-characterized, high-affinity ligand of DARC/ACKR1 (18, 36) and a potent proinflammatory chemokine in the brain that can promote and amplify neuroinflammatory responses through CCR2-dependent signaling (39, 59, 60). MCP-1/CCL2 is strongly induced in the hippocampus following injury, infection, and seizures, conditions associated with robust neuroinflammatory responses (61–63), and has also been implicated in Alzheimer’s disease-associated neuroinflammation (64–66). Consistent with a potential role for MCP-1/CCL2 in the neuroinflammatory phenotype associated with DARC/ACKR1 deficiency, our preliminary observation revealed a marked increase in MCP-1/CCL2 expression throughout the hippocampus of DARC/ACKR1-deficient mice. Loss of DARC/ACKR1-mediated sequestration of MCP-1/CCL2 could increase its local availability and thereby enhance CCR2-dependent signaling, potentially promoting glial responses and the production of additional inflammatory mediators, including MCP-1/CCL2 itself. Such a mechanism could establish a feed-forward inflammatory loop that sustains or amplifies neuroinflammation.

The pronounced neuroinflammatory phenotype observed in DARC/ACKR1-deficient mice was accompanied by significant cognitive deficits, including impaired spatial working memory and object recognition memory. These deficits were also associated with alterations in hippocampal neuronal architecture. DARC/ACKR1-deficient mice exhibited reduced neuronal cell density in the CA1 region, accompanied by thinning of the CA1 pyramidal cell layer. Although less pronounced, the CA2/3 regions also showed a trend toward reduced neuronal cell density and cell-layer thickness. Given the critical roles of these hippocampal subregions in spatial and recognition memory (61, 62), these structural alterations may contribute to the cognitive deficits observed in DARC/ACKR1-deficient mice. The mechanisms underlying these neuronal alterations remain to be determined but may involve both direct effects of altered chemokine signaling on neurons and indirect effects mediated by enhanced neuroinflammation. Loss of DARC/ACKR1 could increase the availability of specific chemokines, thereby enhancing signaling through cognate classical chemokine receptors expressed by neurons. In vitro studies have reported that prolonged exposure of hippocampal neurons to MCP-1/CCL2 reduces neuronal viability, potentially through mechanisms involving glutamatergic excitotoxicity (63). Consistent with these findings, a study of epilepsy reported that increased MCP-1/CCL2/CCR2 signaling was associated with hippocampal neuronal degeneration, whereas genetic deletion of either MCP-1/CCL2 or CCR2 attenuated neuronal loss in the CA3 region (64). In addition, loss of neuronal DARC/ACKR1 may increase the availability of inflammatory chemokines to neighboring glial cells, thereby promoting glial responses and amplifying chemokine-mediated neuroinflammatory signaling. This altered inflammatory environment could, in turn, increase neuronal vulnerability and contribute to neuronal dysfunction and structural alterations.

Our demonstration of neuronal DARC/ACKR1 expression in the human brain suggests that DARC/ACKR1 may similarly contribute to the regulation of neuroinflammatory responses and cognitive function in humans. Human DARC/ACKR1 is polymorphic, with two major codominant alleles, *FYA* and *FYB*, which encode the Fya and Fyb antigens, respectively (65, 66). Fya and Fyb differ by a single amino acid at position 42 in the N-terminal extracellular domain of DARC/ACKR1, with glycine in Fya and aspartic acid in Fyb (65, 66). Although this polymorphism defines distinct Duffy antigens (9, 10), whether the resulting amino acid substitution alters DARC/ACKR1 expression, receptor trafficking, or its chemokine-binding properties remains unclear. Importantly, it is unknown whether Fya and Fyb differ in their expression in neurons, their capacity to sequester chemokines and regulate chemokine availability and distribution in the brain.

Collectively, our findings suggest that neuronal DARC/ACKR1 may function as an endogenous brake on excessive chemokine signaling, thereby limiting neuroinflammation and preserving neuronal and cognitive function. Loss of DARC/ACKR1 may disrupt neuronal regulation of the local chemokine environment, leading to enhanced chemokine signaling and neuroinflammatory responses. The resulting glial responses may further amplify the inflammatory milieu and increase neuronal vulnerability, ultimately contributing to neuronal degeneration and cognitive dysfunction. Further studies are needed to distinguish the direct effects of altered neuronal chemokine signaling from the secondary consequences of neuroinflammation and to define their relative contributions to neuronal and cognitive dysfunction.

## Acknowledgement

We are grateful to the Neuropathology Core at the Emory Alzheimer’s Disease Research Center for providing human postmortem brain tissues. We thank Ning Wu and Shu Chen for technical support.

---

**Supplementary Figure 1.**
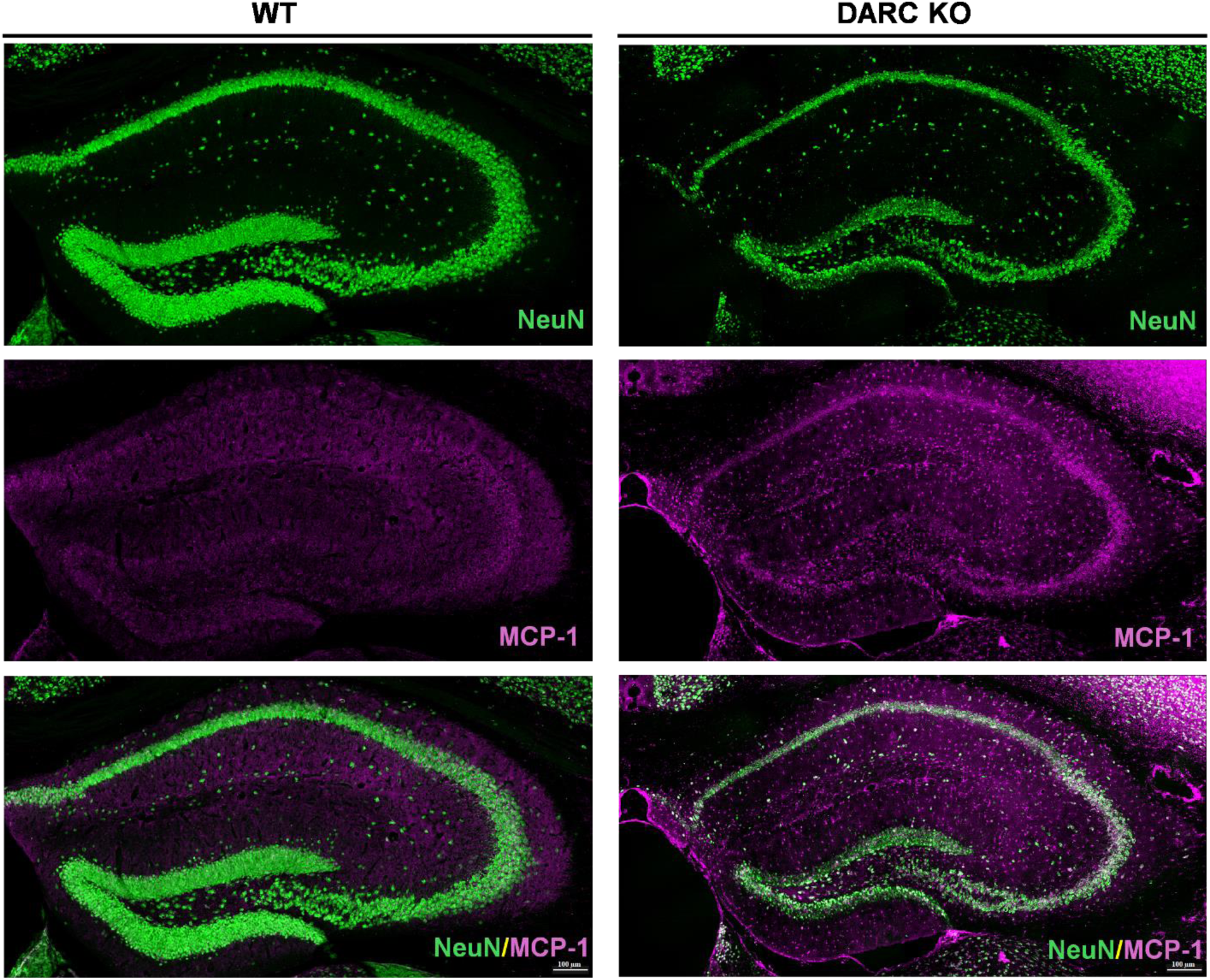
DARC/ACKR1 deficiency increases MCP-1/CCL2 levels in the hippocampus. Representative confocal images of the hippocampus from wild-type (WT) and DARC/ACKR1-deficient (DARC KO) mice immunostained for NeuN (green, neurons) and MCP-1/CCL2 (magenta). Top row, NeuN; middle row, MCP-1/CCL2; bottom row, merged images.

